# Astrocyte extracellular matrix modulates neuronal dendritic development

**DOI:** 10.1101/2024.08.06.606424

**Authors:** Joel G. Hashimoto, Nicholas Margolies, Xiaolu Zhang, Joshua Karpf, Yuefan Song, Brett A. Davis, Fuming Zhang, Robert J. Linhardt, Lucia Carbone, Marina Guizzetti

## Abstract

Major developmental events occurring in the hippocampus during the third trimester of human gestation and neonatally in altricial rodents include rapid and synchronized dendritic arborization and astrocyte proliferation and maturation. We tested the hypothesis that signals sent by developing astrocytes to developing neurons modulate dendritic development *in vivo*. We altered neuronal development by neonatal (third trimester-equivalent) ethanol exposure in mice; this treatment increased dendritic arborization in hippocampal pyramidal neurons. We next assessed concurrent changes in the mouse astrocyte translatome by translating ribosomal affinity purification (TRAP)-seq. We followed up on ethanol-inhibition of astrocyte *Chpf2* and *Chsy1* gene translation because these genes encode for biosynthetic enzymes of chondroitin sulfate glycosaminoglycan (CS-GAG) chains (extracellular matrix components that inhibit neuronal development and plasticity) and have not been explored before for their roles in dendritic arborization. We report that *Chpf2* and *Chsy1* are enriched in astrocytes and their translation is inhibited by ethanol, which also reduces the levels of CS-GAGs measured by Liquid Chromatography/Mass Spectrometry. Finally, astrocyte-conditioned medium derived from *Chfp2*-silenced astrocytes increased neurite branching of hippocampal neurons *in vitro*. These results demonstrate that CS-GAG biosynthetic enzymes in astrocytes regulates dendritic arborization in developing neurons.

## Introduction

Neurodevelopment is highly dynamic and ontogenic processes are tightly regulated in their inception and termination (1, 2). The neonatal period between birth and post-natal day (PD) 7 (mice) or PD 9 (rats) corresponds to the third trimester of human gestation with regards to brain development and is characterized by the rapid and synchronized growth of the dendritic arbor of pyramidal neurons and the proliferation and maturation of astrocytes in the cortex and hippocampus (3, 4). Astrocytes have been identified as playing key roles in neurodevelopment and neurodevelopmental disorders (4-7). However, the understanding of the contribution of astrocytes to dendritic development *in vivo* is limited.

To test the hypothesis that immature astrocytes send signals to immature neurons that modulate the development of the dendrites during the first postnatal week in mice *in vivo*, we altered the normal development of the dendritic arbor by exposing mice to a developmental neurotoxicant, ethanol, during the first post-natal week. The mouse model used in this study mimics prenatal alcohol exposure (PE) during the third trimester-equivalent of human gestation. In humans, prenatal ethanol exposure (PE) causes Fetal Alcohol Spectrum Disorders (FASDs), a cause of lifelong functional disabilities including cognitive and behavioral impairment associated with psychiatric comorbidities (8). The prevalence of FASD is estimated to be 1-5% in the United States (9-11), though FASD prevalence can be 10-to 40-times higher in certain subpopulations (12). While FASDs are estimated to be as prevalent as autism spectrum disorder, they are often under- or mis-diagnosed (8). Clinical and preclinical studies indicate that neuronal plasticity and connectivity in the hippocampus are affected by *in utero* ethanol exposure, and that these alterations may play a role in FASD central nervous system dysfunction (13-15). Children with PE have deficits in hippocampus-mediated processes (16, 17) and altered hippocampal connectivity has been reported in humans and animal models (18-20). Preclinical studies have also identified alterations in hippocampus-dependent behaviors following developmental ethanol exposure (21-23). Here we found that the exposure to ethanol vapor between PD2 and PD7 increased the arborization of CA1 pyramidal neurons.

We next characterized perturbations in the astrocyte translatome by ethanol vapor using the translating ribosomal affinity purification (TRAP) approach in Aldh1l1-EGFP/Rpl10a mice (24, 25) to identify mechanistic leads on astrocyte factors that modulate the increase in dendritic development observed after ethanol exposure. We characterized changes in gene translation related to functions that are both astrocyte-specific as well as non-astrocyte-specific but altered selectively in astrocytes.

Chondroitin sulfate glycosaminoglycans (CS-GAGs) are major components of the brain extracellular matrix (ECM). These sugar chains are inhibitors of neuronal development and are key to proper dendrite and axonal growth and arborization and are also key components of perineuronal nets, which restrict plasticity and stabilize synapses, though at the developmental stage investigated in this study perineuronal nets have not developed yet (26). From the translatome analysis we found that translation of *Chpf2* and *Chsy1*, two enzymes involved in the biosynthesis of CS-GAGs, are downregulated by ethanol exposure in astrocytes. Following up on this observation, we found that neonatal exposure decreased hippocampal CS-GAG disaccharide levels and that hippocampal pyramidal neurons cultured with astrocyte-conditioned medium (ACM) prepared from astrocyte cultures in which *Chpf2* was silenced extended longer minor neurites and increased the branching and complexity of the neurites.

In conclusion, we identified an astrocyte-mediated mechanism of dendritic development of CA1 pyramidal neurons involving the regulation of CS-GAG biosynthetic enzymes. Alteration in the expression of these enzymes may play a role in the disruption of proper dendritic development in FASD and, possibly, other neurodevelopmental disorders.

## Results

### Neonatal ethanol exposure increases dendritic arborization of mouse CA1 pyramidal hippocampal neurons

To investigate astrocyte-mediated mechanisms of dendrite development, we altered normal dendritic development of CA1 pyramidal neurons with ethanol. We find that neonatal ethanol exposure increased complexity, the sum of terminal orders, the length, the number of ends and the number of nodes in apical dendrites determined by Golgi-Cox staining followed by morphometric analysis (**Figure 1**). In basal dendrites, ethanol increased complexity, the maximal terminal distance, the sum of terminal orders, the total basal dendrite length, the average basal dendrite length, the number of ends and the number of nodes (**Figure 2**). Representative neuron tracings from male and female PD7 mice after air (control) or ethanol vapor exposure are shown in **Figure 1A**.

**Figure 1.**
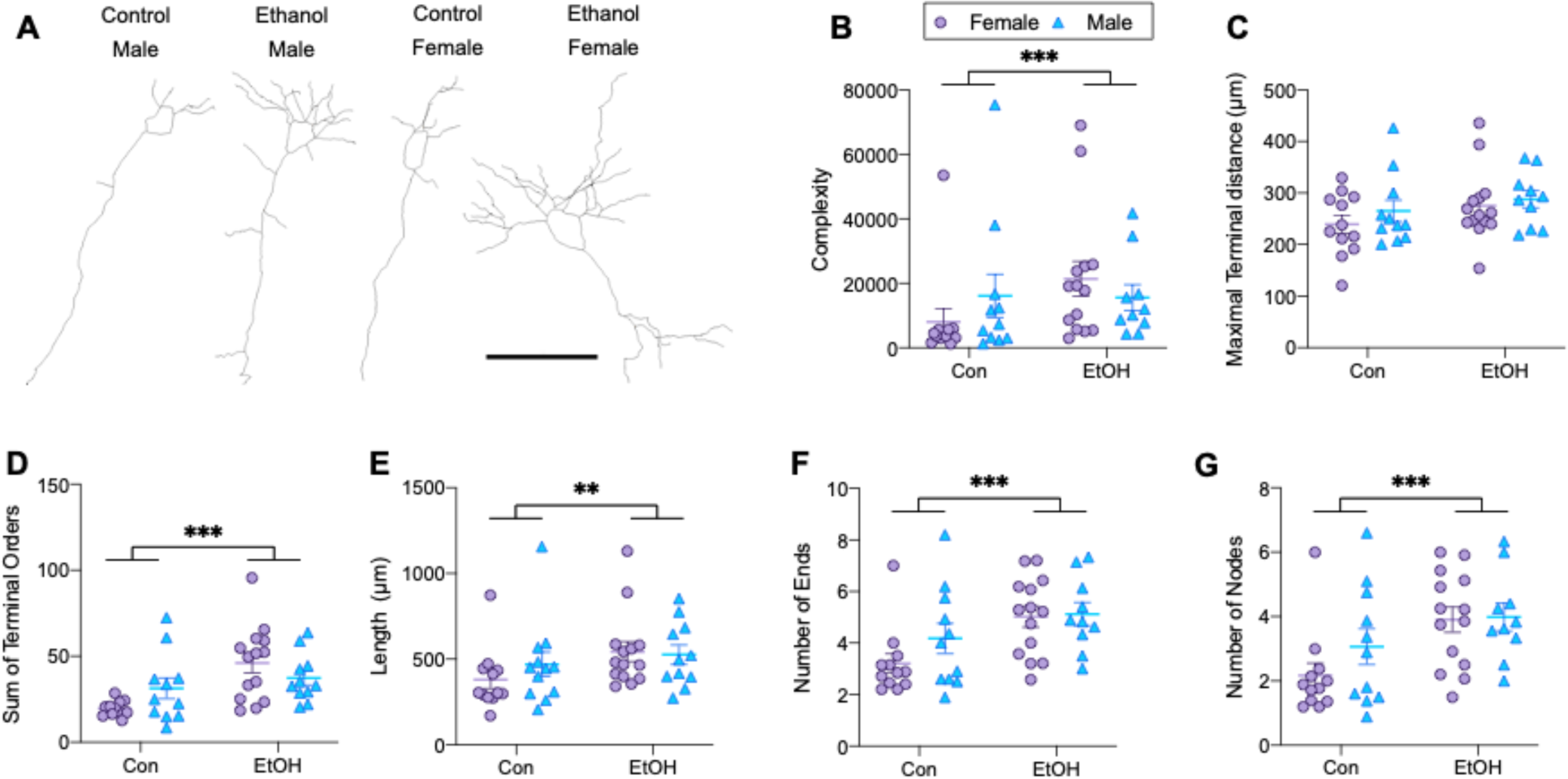
Morphometric analysis of CA1 pyramidal neuron apical dendrites from PD7 mice neonatally exposed to ethanol vapor or air control. **A**: Representative Neurolucida tracings of PD7 CA1 pyramidal neurons (scale bar 50 µm). **B**: Ethanol exposure increased complexity (p = 0.0004). **C**: Maximal terminal distance was not altered by ethanol exposure. The sum of terminal orders (**D**; p = 5.0x10^-5^), the length of the apical dendrite (**E**; p = 0.002), the number of ends (**F**; p = 0.0006), and the number of nodes (**G**; p = 0.0007) of the apical dendrite were increased by ethanol exposure.

**Figure 2.**
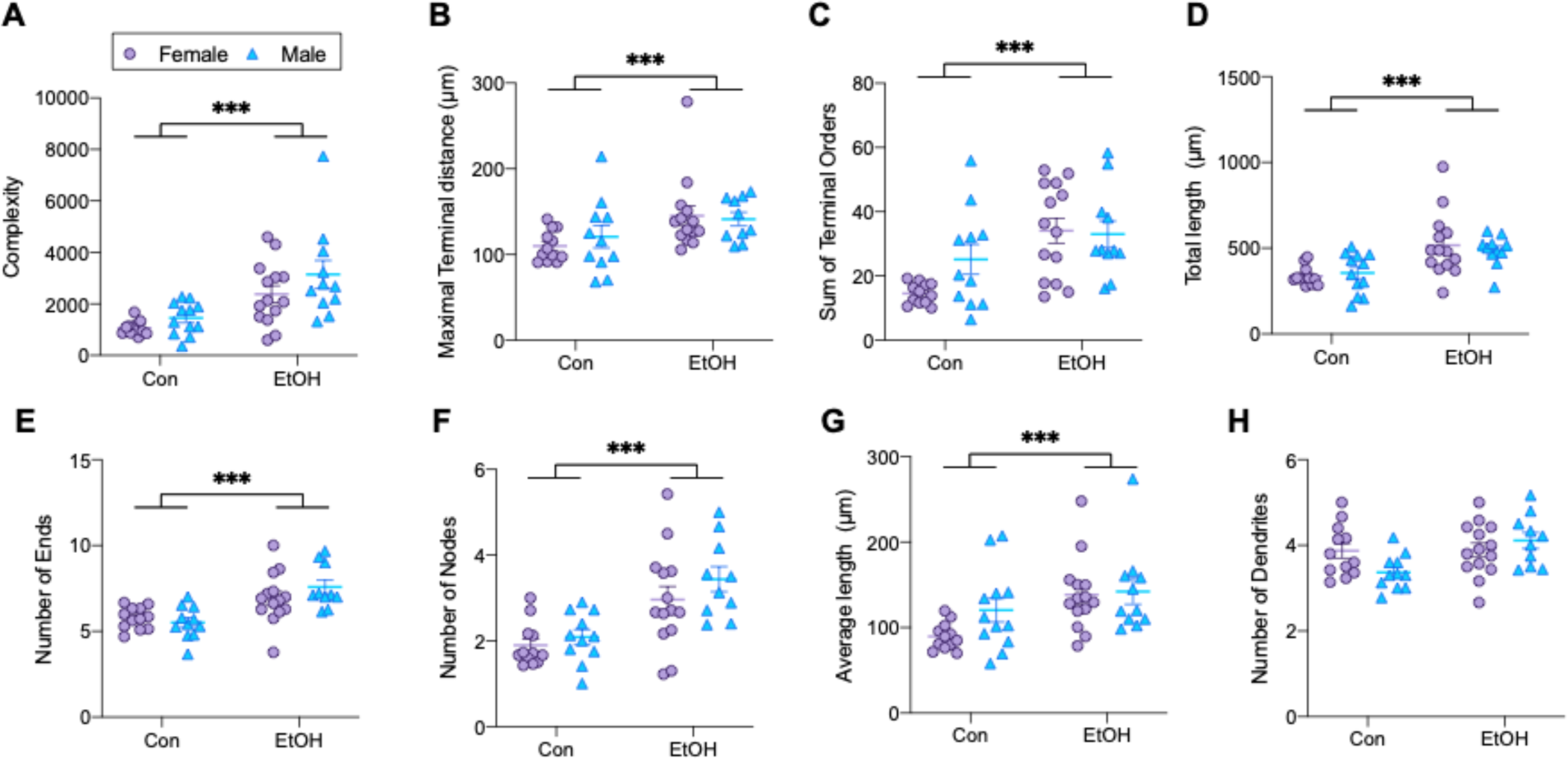
Morphometric analysis of CA1 pyramidal neuron basal dendrites from PD7 mice neonatally exposed to ethanol vapor or air control. Ethanol exposure increased basal dendrite complexity (**A**; p = 8.4 x 10^-6^), the maximal terminal distance (**B**; p = 0.0006), the sum of terminal orders (**C**; p = 4.2 x 10^-5^), the total dendrite length (**D**; p = 0.0006), the number of ends (**E**; p = 6.7 x 10^-5^); the number of nodes (**F**; p = 0.0003) and the average dendrite length (**G**; p = 5.9 x 10^-5^). **H**: The number of basal dendrites was not altered by ethanol exposure.

At this developmental stage dendritic spines are sparse and immature (27); dendritic spine density in secondary branches of apical dendrites was not affected by ethanol exposure when spine density was normalized to the length of the dendrite segment (**Figure 3A**). However, spine density was increased in the ethanol exposed animals relative to controls when normalized to both length and diameter (**Figure 3B**); as the diameters of the dendrites were smaller in ethanol exposed animals (**Figure 3C**). Representative 100x images of dendrite segments showing dendrite thickness and dendritic spines are show in **Figure 3D**. Together, these results indicate that neonatal ethanol increased dendritic arborization and induced the thinning of the dendrites.

**Figure 3.**
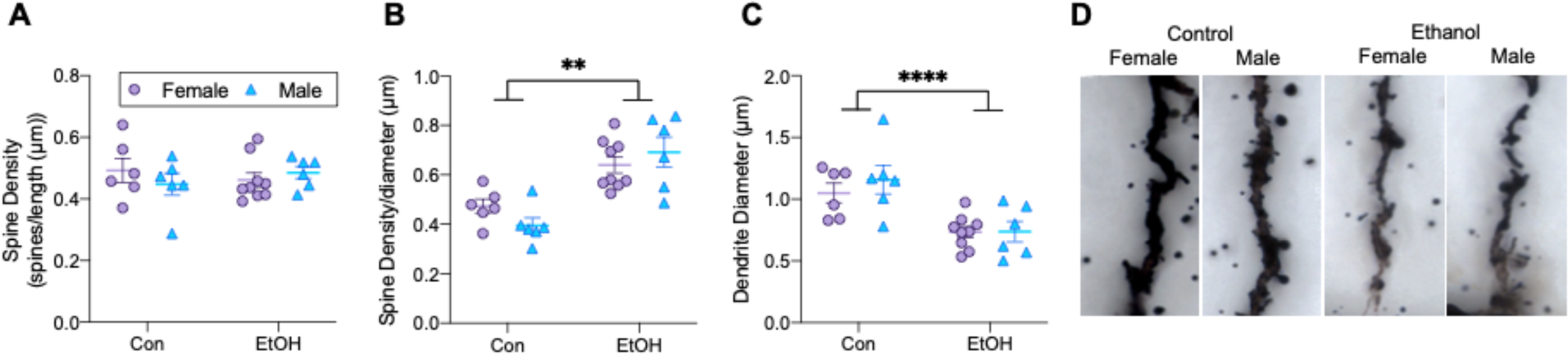
Analysis of spine density and dendrite diameter in secondary branches of apical dendrites following neonatal ethanol exposure. **A**: The number of spines per length of the dendrite (µm) was not altered by ethanol. **B**: The number of spines per length (µm) per diameter (µm) of the dendrite was increased by neonatal ethanol exposure (p = 0.003). **C**: Ethanol exposure decreased the diameter of the secondary branches of the apical dendrite from which spine density was determined (p = 9.5 x 10^-5^). **D**: Representative images of Golgi-Cox-stained dendrites analyzed for spine density and dendrite diameter at 100x magnification.

### Effect of neonatal ethanol exposure on the astrocyte translatome

Aldh1l1-EGFP/Rpl10a mice expressing an EGFP-Rpl10a ribosomal transgene targeted to cells expressing *Aldh1l1* (a specific astrocyte marker) was used to evaluate changes in astrocyte ribosome-associated actively translating RNA by the TRAP method (24, 25). The TRAP procedure on the hippocampus from PD7 Aldh1l1-EGFP/Rpl10a mice resulted in a 6- to 7-fold enrichment in RNA levels of astrocytic markers and 3- to 6-fold depletion in RNA levels of neuronal, microglia, and oligodendrocyte markers, when compared to the input by qRT-PCR (**Figure 4A**).

**Figure 4.**
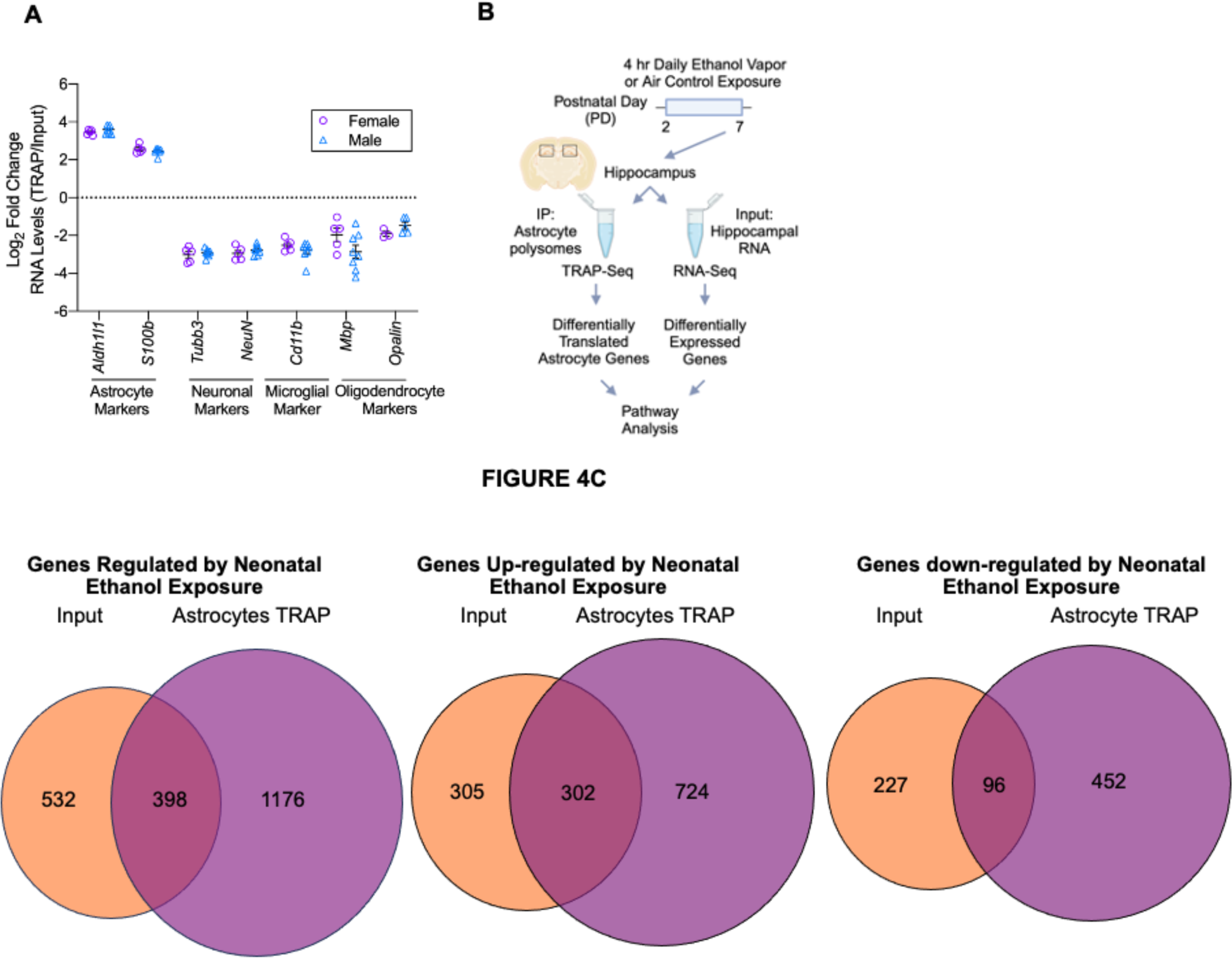
**A**: qRT-PCR of cell-type marker genes in the TRAP fractions vs input fractions shows high enrichment of astrocyte markers and depletion of neuronal, microglial, and oligodendrocyte markers. **B**: Schematic representation of the design of astrocyte TRAP-seq experiments. **C:** Venn diagrams comparing ethanol regulated (left), ethanol upregulated (center), and ethanol downregulated (right) genes in TRAP and input samples and their overlap.

The hippocampi from 12 control pups (6 males and 6 females) and 12 pups exposed to ethanol vapor (6 males and 6 females), one male and one female pup per litter, underwent the TRAP procedure followed by RNA sequencing (TRAP-seq). We additionally sequenced the respective input samples, consisting of aliquots of RNA taken before TRAP pull-down (RNA-seq). (**Figure 4B**; schematic of experimental design). There were no significant differences in body weights of PD7 male and female mice or between air control and ethanol exposed mice, and there were no significant differences in BACs between male and female mice (**Table 1**).

**Table 1.**
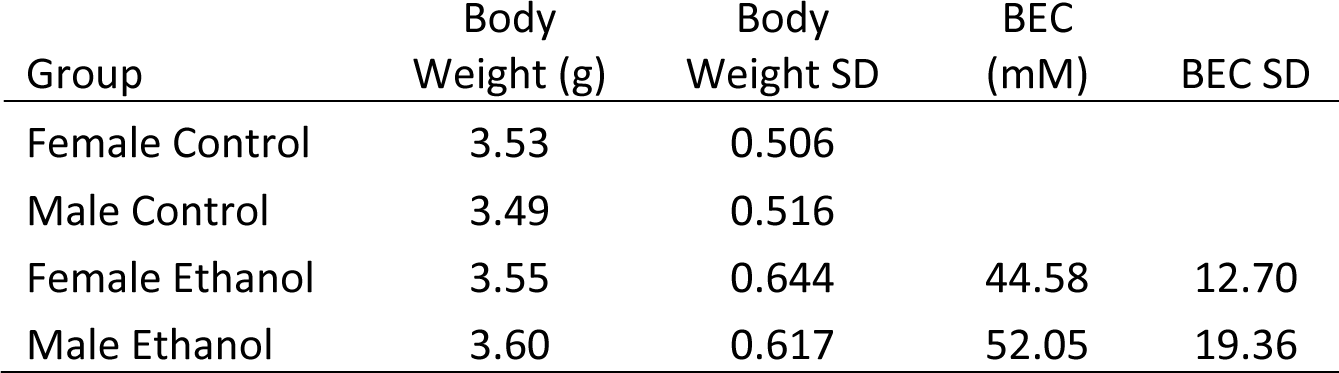
Female and male mouse pups were weighed on PD7, after the last ethanol vapor or air exposure; Blood Alcohol Concentrations (BACs) from trunk blood were determined by gas chromatography. SD: standard deviation.

The RNA from TRAP and input samples had RNA Integrity Numbers (RIN) ranging between 8.8 and 10 (**Suppl. Table 1**). **Suppl. Figure 1** shows that RNA yield after pull-down (**A**), EGFP astrocyte translation (**B**) and expression (input; **C**), and Rpl10a normalized counts from TRAP-seq (**D**) and RNA-seq (**E**) analyses did not differ across sexes or treatments indicating that the TRAP method is not biased by sex or neonatal ethanol treatments.

The comparison of the TRAP-seq results from the hippocampi of ethanol vapor-exposed pups vs air-exposed pups resulted in 1,574 astrocyte differentially translated (DT) genes, of which 1,026 genes (65%) were upregulated and 548 genes (35%) were downregulated. In the input fraction 932 genes were differentially expressed (DE); 607 of which (65%) were upregulated, and 323 (35%) were downregulated. Moreover, we found that 303 genes were upregulated in both TRAP and input samples, 96 genes were downregulated in both TRAP and input samples. Venn diagrams of genes regulated (left), upregulated (center) and downregulated (right) by ethanol vapor in TRAP and input samples and their overlap are shown in **Figure 4C**. The complete lists of DT and DE genes are shown in **Suppl. Tables 2-3**.

DE and DT genes were analyzed for overrepresentation in gene ontology (GO) and known pathways. The top 30 categories significantly overrepresented in EnrichR pathway analysis of DT genes data were dominated by terms related to ribosomal biogenesis and function and protein translation (**Suppl. Figure 2A**). In contrast, the top 30 categories of DE genes included several signal transduction pathways (**Suppl. Figure 2B**). Interestingly, there was no overlap between the top 30 overrepresented categories in the TRAP and in the input data sets.

We next ran pathway analysis separately on upregulated (**Suppl. Figure 3A**) and downregulated (**Suppl. Figure 3B**) DT genes. Top significant pathways in DT genes upregulated by ethanol include “axon guidance” and “neuronal projection”. Genes in these pathways include ephrin receptor B6 (*Ephb6*), a transmembrane protein involved in cell adhesion and neurite outgrowth, *Slit1*, encoding for a secreted protein that induces axonal and dendritic extension and provides axonal and dendritic guidance cues, and *Col41A*. Upregulation of these pathways, and these genes in particular, is consistent with increased dendritic arborization induced by ethanol. Additional significant categories in the analysis of the genes upregulated by ethanol include terms involved in synaptic function (**Suppl. Figure 3A**). While synaptogenesis is only beginning at this developmental stage, it is well-established that astrocytes express a wide array of neurotransmitter receptors and voltage-gated ion channels that contribute to synapse development and excitability in neurons (28-30). Pathways identified for DT downregulated genes include ribosome biogenesis and protein translation (**Suppl. Figure 3B).**

### Pathway analyses of astrocyte DT genes enriched, and non-enriched in astrocytes

We next identified DT genes that were astrocyte-enriched or not astrocyte-enriched; we defined astrocyte enrichment as genes with higher RNA reads in the TRAP-seq samples than in the RNA-seq (input) samples (adjusted p <0.05 and a log_2_ fold change ≥ 1). It is assumed that DT genes enriched in TRAP-seq samples are associated with astrocyte-specific functions, while DT genes that are non-astrocyte enriched are associated with functions also present in other cell types but regulated by ethanol in astrocytes. As shown in **Table 2**, 1,301 DT genes were non-astrocyte enriched (874 of these genes were upregulated and 427 downregulated). Pathway analysis of non-astrocyte-enriched DT genes included numerous categories related to protein translation and ribosome biogenesis and function (**Figure 5A**); most of these categories were also overrepresented in the analysis of down-regulated genes (**Figure 5C**). Pathway analysis of non-astrocyte-enriched upregulated genes identified several categories involved in the synaptic excitability of neurons, including neurotransmitter receptors and voltage-dependent ion channels (known to be expressed both in neurons and astrocytes), and categories related to the positive regulation of dendritic extension and neuronal processes (**Figure 5B**). We marked with green arrows pathways related to ribosome biogenesis and protein translation; with red arrows pathways related to neuronal projection and dendrite extension; and with tan arrows pathways related to synaptic function and excitability of neurons (**Figure 5A-C**). Pathway analyses of non-astrocyte-specific genes shown in **Figure 5** largely overlap with pathway analyses of the entire ethanol regulated genes (**Suppl Figure 2A**; **Suppl. Figure 3A-C**), consistent with the fact that most of the regulated genes are not astrocyte-specific (see **Table 2**).

**Figure 5.**
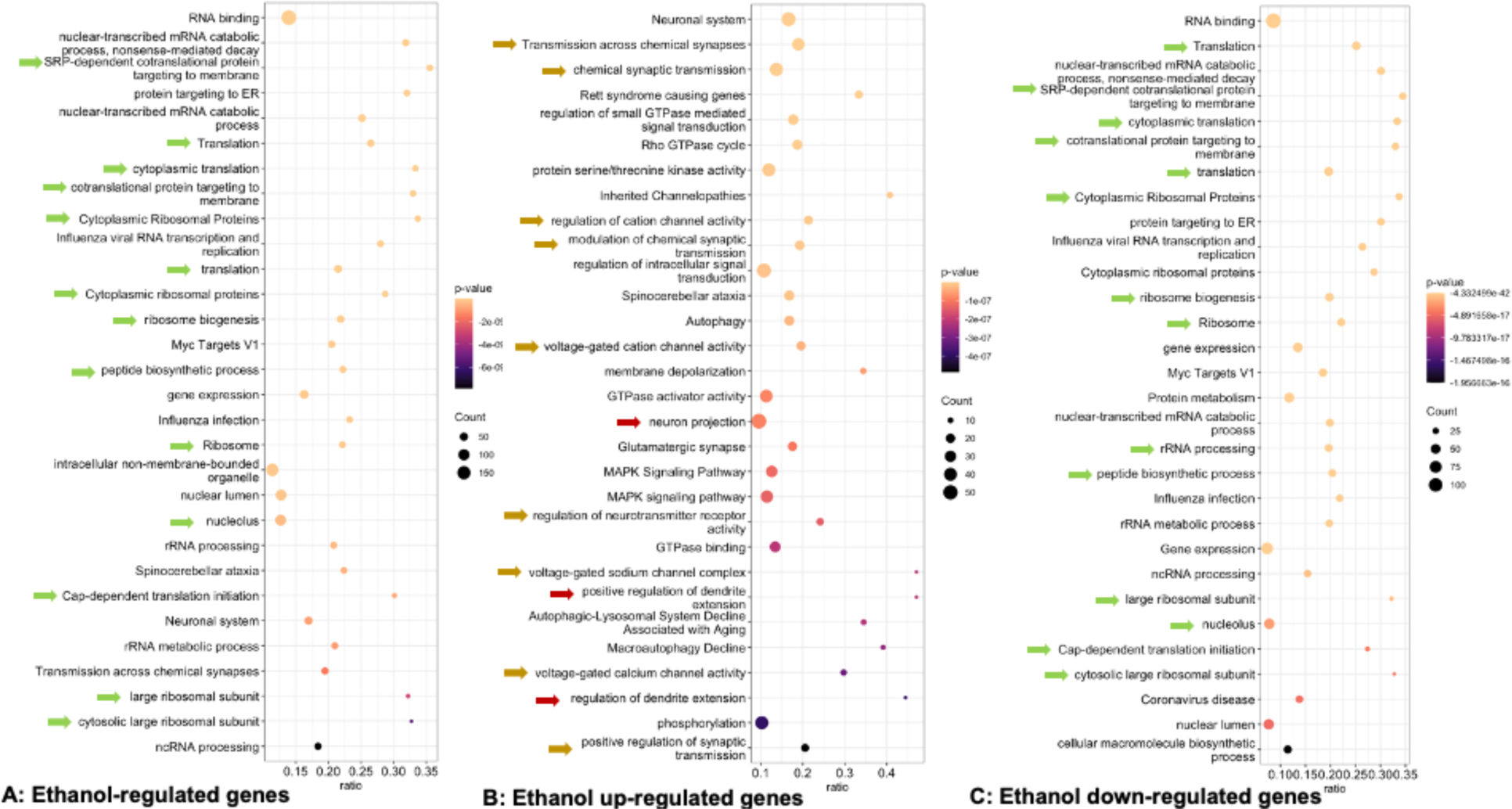
Category overrepresentation summary plots of ethanol regulated genes in the TRAP (astrocyte) fraction of genes that were not enriched in astrocytes. **A**: Ethanol regulated genes not enriched in astrocytes analyzed for Gene Ontology (GO), and pathway overrepresentation using EnrichR. The top 30 gene categories based on p-values are shown. In each plot, the size of the dot represents the number of genes that were significantly regulated, and the color scale represents the p-value, with lighter colors indicating lower p-values. Categories marked with green arrows are related to gene translation. **B**: Top overrepresented categories derived from GO analysis of genes upregulated by ethanol and not enriched in astrocytes. Gene categories with tan arrows relate to synaptic activity and voltage-gated channels and red arrows relate to dendrite growth. **C**: Top overrepresented categories derived from GO analysis of genes downregulated by ethanol and not enriched in astrocytes. Gene categories marked with green arrows relate to gene translation.

**Table 2:** Number of genes differentially translated after neonatal alcohol exposure.

| Groups of ethanol-regulated genes | Number of ethanol-regulated genes | Number of genes upregulated by ethanol | Number of genes downregulated by ethanol |
| --- | --- | --- | --- |
| Enriched in Astrocytes | 273 | 152 | 121 |
| Not Enriched in Astrocytes | 1301 | 874 | 427 |
| Total | 1574 | 1026 | 548 |

Neonatal ethanol altered the translating RNA of 273 astrocyte-enriched genes (152 of these genes were upregulated and 121 downregulated by ethanol) (**Table 2**). **Figure 6** shows top categories of astrocyte-enriched genes regulated (**A**), upregulated (**B**), and downregulated (**C**) by ethanol. Pathway analysis of astrocyte-enriched ethanol upregulated DT genes identified terms involved in cholesterol biosynthesis/cholesterol metabolism (blue arrows) and in cell adhesion/axon guidance (magenta arrows) among the top significantly overrepresented categories. Among the top significantly overrepresented terms in pathway analysis of differentially translated genes that are enriched in astrocytes and downregulated by ethanol were terms involving CS-GAG biosynthesis (**Figure 6C**; purple arrows). Complete pathway analyses are provided in **Suppl. Table 4**.

**Figure 6.**
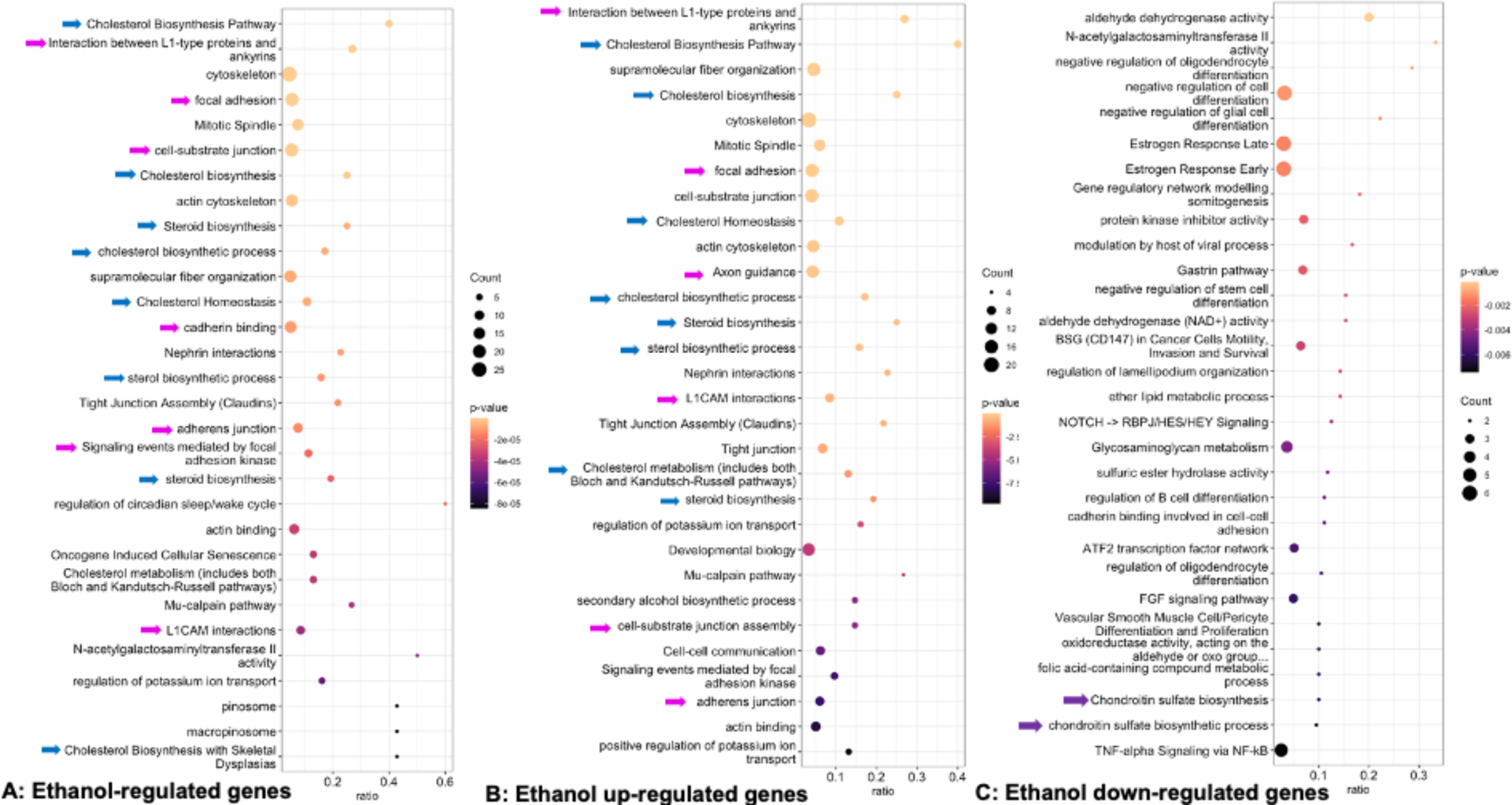
Category overrepresentation summary plots of ethanol regulated genes in the TRAP (astrocyte) fraction of genes that were enriched in astrocytes. **A**. Top overrepresented categories derived from GO analysis of ethanol-regulated genes enriched in astrocytes. Categories marked with blue arrows relate to steroid biosynthesis and with magenta arrows to cell-cell interactions, cell-ECM interactions, and cell adhesion. **B**: Top overrepresented categories derived from GO analysis of genes upregulated by ethanol and enriched in astrocytes analyzed with EnrichR. Categories with blue arrows relate to steroid biosynthesis and magenta arrows relate to cell-cell interactions, cell-ECM interactions, and cell adhesion. **C**: Top overrepresented categories derived from GO analysis of genes downregulated by ethanol and enriched in astrocytes analyzed with EnrichR. Gene categories with purple arrows are related to chondroitin sulfate biosynthesis.

From the analyses in **Figures 5** and **6** and **Suppl. Figure 3** it emerges that significantly overrepresented pathways of upregulated genes include “axon guidance”, “neuron projection”, “positive regulation of dendrite extension”, “focal adhesion”, and “L1CAM interaction” all of which are involved in cell adhesion (necessary for the growth of neuronal processes) and axon and dendrite development while significantly overrepresented pathways of downregulated genes include “chondroitin sulfate biosynthesis” and “chondroitin sulfate biosynthetic process” involved in the inhibition of neuronal development and plasticity (see next paragraph) which strongly supports the increased dendritic arborization observed in **Figures 1-2**.

### Neonatal ethanol exposure decreases levels of CS-GAG disaccharides in the rat hippocampus

The overrepresented categories “Chondroitin sulfate biosynthesis” and “Chondroitin sulfate biosynthetic processes” marked by a purple arrow in the analysis of astrocyte-enriched genes downregulated by ethanol (**Figure 6C**) had 2 genes, *Chpf2* and *Chsy1*, encoding for biosynthetic enzymes specific to chondroitin sulfate (CS) chains (31). CS-GAG chains consist of repeating disaccharides covalently attached to proteoglycans (PGs) and are inhibitiors of neuronal process development and plasticity (26). Quantitative RT-PCR confirmed that neonatal ethanol exposure downregulates *Chpf2* and *Chsy1* translation in astrocytes (**Figure 7A**). We also confirmed *Chpf2* and *Chsy1* expression in astrocytes by RNA-FISH (**Figure 7B**). *Chpf2* and *Chsy1* are about 2- and 4-fold enriched in astrocytes, respectively (**Figure 7C)**. In agreement with the downregulation of *Chpf2* and *Chsy1* by ethanol, we found decreased levels of total CS-GAG disaccharides, as well as decreased levels of chondroitin-4-sulfate (C4S)-, C6S-, and C0S-GAGs (which together represent about 99% of all total CS-GAGs in the neonatal hippocampus; **Suppl. Figure 4**) measured by LC/MS in the hippocampus of neonatal rats after ethanol exposure (**Figure 7D-G**).

**Figure 7.**
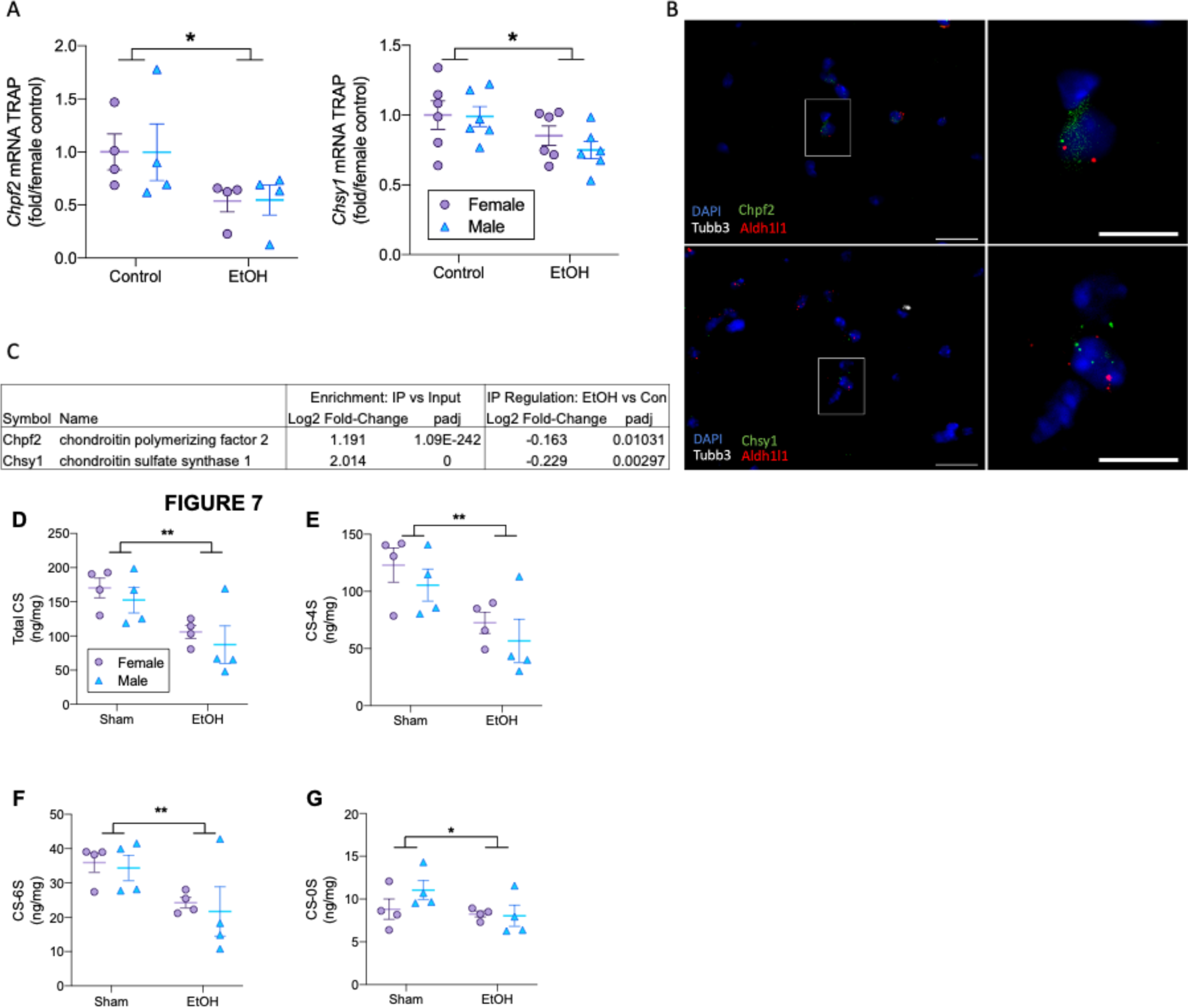
Neonatal ethanol exposure decreases chondroitin sulfate glycosaminoglycan (CS-GAG) biosynthetic enzymes and CS-GAGs levels in the neonatal hippocampus. **A**: Down-regulation of *Chpf2* (left) and *Chsy1* (right) in the TRAP fraction by qRT-PCR (*Chpf2* p-value = 0.027; *Chsy1* p-value = 0.022). **B**: *Chpf2* (top panels) and *Chsy1* (bottom panels) expression in astrocytes of the CA1 region of the hippocampus by RNA-FISH. Left panels: 100x objective images of RNA-FISH probes for *Chpf2* (green, top) and *Chsy1* (green, bottom) and cell-type markers *Aldh1l1* (red, astrocytes) and *Tubb3* (white, neurons); nuclei are stained by DAPI (Blue; scale bar = 20 µm). Right panels: zoomed region inside the white box of the left panels (scale bar = 10 µm). Staining puncta show proximal presence of *Aldh1l1* and *Chpf2* (top panels) or *Aldh1l1* and *Chsy1* (bottom panels) staining. **C**: Astrocyte enrichment and ethanol regulation of *Chpf2* and *Chsy1* in TRAP-Seq data. Concentrations (ng/mg tissue) of total CS-GAG disaccharides (**D**; p = 0.001), CS-4S disaccharides (**E**; p = 0.002), CS-6S disaccharides (**F**: p = 0.005), and CS-0S disaccharides (**G**; p = 0.01) were decreased following ethanol treatment in the neonatal rat hippocampus. CS-GAG disaccharides were quantified by LC/MS and normalized to hippocampus weight.

### Hippocampal neuron neurite outgrowth is upregulated by ACM prepared from Chpf2-silenced astrocyte cultures

Finally, we tested whether the silencing of *Chfp2* in astrocytes was able to inhibit neurite outgrowth in hippocampal neurons *in vitro*. ACM prepared from astrocytes in which *Chpf2* expression was silenced increased the complexity, the number of nodes and ends, and the average minor neurite length (i.e., the nascent dendrites) of hippocampal neurons in culture compared to neurons exposed to ACM prepared from control astrocytes (**Figure 8 A-D**). **Figure 8E** shows representative neurons in the two experimental conditions. The treatment of astrocytes with a specific *Chpf2* SiRNA resulted in a decrease in *Chfp2* expression by 95.3% (SEM 0.11%) as determined by qRT-PCR.

**Figure 8.**
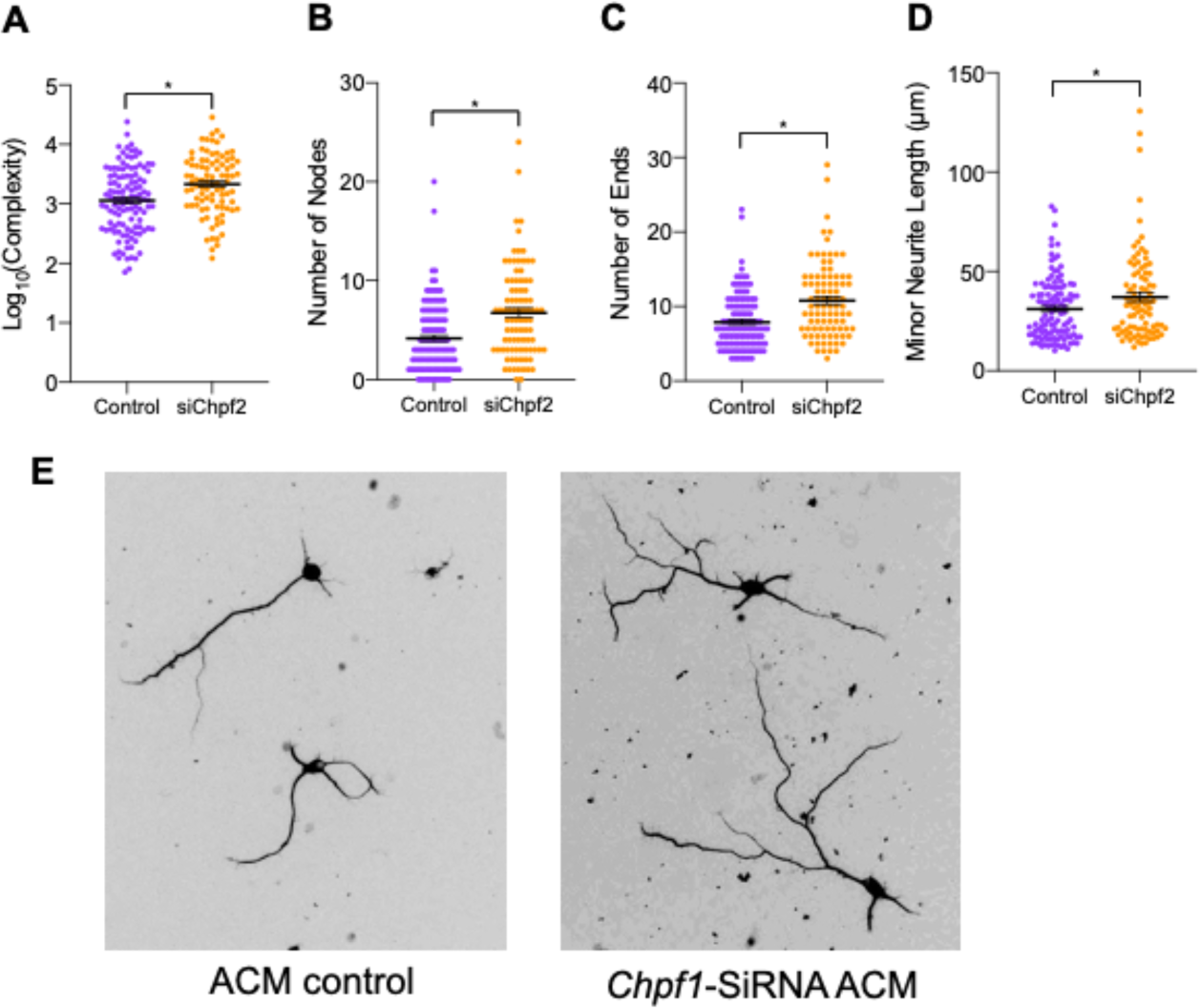
ACM from *Chpf2* siRNA-treated astrocytes increases neurite outgrowth in hippocampal neurons *in vitro*. Primary astrocyte cultures were treated with *Chpf2* SiRNA or control; ACM was collected and transferred to primary cortical neuron cultures for 48 h. Hippocampal pyramidal neurons incubated in the presence of ACM from astrocytes in which *Chpf2* was silenced showed increased neuronal neurite complexity (A; p = 0.03); number of nodes (**B**; p = 0.05), number of ends (**C**; p = 0.01), and the length of the minor neurites (**D**; p = 0.03). **E**: representative neurons exposed to control ACM (left) or ACM prepared from astrocytes whose *Chpf2* expression was silenced.

## Discussion

To identify astrocyte-mediated mechanisms regulating dendritic arborization we altered the process of dendritic arborization at a time point in development when the rapid and synchronized increase in the neuronal dendritic arbor and astrocyte proliferation and maturation are occurring (1-3). The FASD model used in this study allowed us to characterize changes in astrocyte translatome during this narrow developmental window and link them to increased dendritic arborization.

In agreement with the translatome results, we found increased dendritic arborization of apical and basal dendritic arbors of CA1 pyramidal neurons in neonatal mice exposed to ethanol vapor (**Figures 1**,**2**). This dysregulation of neuronal development that alters the developmental trajectories was further reflected by decreased apical dendrite diameters and increased spine densities per dendritic area (**Figure 3**) following ethanol vapor exposure. During this developmental stage of rapid and synchronized growth of the dendritic arbor in pyramidal neurons of the cortex and hippocampus, alterations in both directions (i.e., hypoconnectivity and hyperconnectivity) of this programming lead to altered local and distal neuronal connectivity associated with functional and behavioral dysfunction (32). We and others have previously reported that neonatal or prenatal ethanol exposure increases dendritic arborization in rats (33-35).

We employed astrocyte-TRAP-seq in combination with bulk tissue RNA-seq to investigate changes induced selectively in astrocytes by developmental ethanol exposure that could be responsible for altered dendritic arborization. The higher number of genes differentially translated (DT genes) in astrocytes versus the number of genes differentially expressed (DE genes) in the bulk hippocampus demonstrates that the cell-type specific approach of the TRAP procedure enhances the detection of ethanol regulated genes in astrocytes partially obscured in the heterogeneous mixture of cell types in the bulk hippocampus (**Figure 4C**, **Table 2**).

Astrocytes have many functions in the developing brain, some of which are common to other cell types, while others are specific to astrocytes. The sequencing of input samples together with the TRAP samples allowed us to run pathway overrepresentation analyses separately on genes that are enriched in astrocytes (representing astrocyte-specific functions altered by ethanol) and on genes that are not enriched in astrocytes (representing functions that are common to other cell types but are selectively altered by ethanol in astrocytes). This compartmentalization of pathway analyses was instrumental in bringing attention to astrocyte-specific functions altered by ethanol.

Categories overrepresented in pathway analyses of astrocyte-enriched genes upregulated by ethanol included several terms related to cell adhesion (which is essential to dendritic growth) and axon and dendrite development (**Figure 5B**, red arrows; **Figure 6A, B**; magenta arrows); conversely, terms involved in the inhibition of neuronal development are downregulated (**Figure 3C**; purple arrows), consistent with the observed increased in dendritic arborization induced by ethanol. Astrocytes are major producers of brain ECM proteins (36) that can modulate dendritic development and the secretion of ECM proteins is known to be altered by ethanol (37). Astrocytes also express CAMs in their membranes that interact with homologous and heterologous proteins on the neuronal membrane providing the adhesion necessary for the growth of neuronal processes (38).

We followed up on the decreased translation of astrocyte-enriched *Chpf2* and *Chsy1,* the 2 genes included in the statistically overrepresented categories “chondroitin sulfate biosynthesis” and “chondroitin sulfate biosynthetic processes” (**Figure 6C**; purple arrows). *Chpf2* and *Chsy1* encode enzymes involved in the biosynthesis of CS-GAGs, unbranched polysaccharides, consisting of repeating disaccharide units (glucuronic acid and *N*-acetylgalactosamine) modified by sulfation in positions 2-, 4-, and/or 6-, covalently bound to core-protein proteoglycans (PGs) (39, 40). CS-PGs of the lectican family are the most abundant proteins of the CNS ECM (41). Lecticans are highly expressed in the developing brain where they serve as barriers to cell migration and to the growth of neuronal processes and are essential to the formation of the proper neuronal connectivity (26, 38). Astrocytes are the major producers of lecticans brevican and neurocan in the PD7 cortex and hippocampus of Aldh1l1-EFGP-Rpl10a mice (42) (Zhang et al, in preparation). *Chpf2* and *Chsy1* are enriched in astrocytes (consistent with astrocytes being the major source of CS-PG lecticans) and are downregulated by ethanol in TRAP-seq and TRAP-qPCR analyses (**Figures 6C** and **7A,B,C**). We also found that CS-GAG disaccharide levels are reduced by ethanol in the hippocampus of neonatal rats (**Figure 7D-G**). In agreement with these findings, we also report an increase in branching (number of nodes and ends), complexity, and the length of minor neurites (the nascent dendrites) in hippocampal neurons cultured in the presence of ACM derived from astrocyte cultures in which *Chpf2* expression is silenced (**Figure 8**), mechanistically linking ethanol-induced increased dendritic arborization to the downregulation of CS-GAG biosynthetic enzymes in astrocytes. Individuals with FASD also show skeletal abnormalities due to altered bone development (43, 44) and CS-GAGs are involved in bone mineralization (45). Therefore, the down-regulation by ethanol of CS-GAGs may also contribute to these peripheral abnormalities.

Our translatomic results also provided several new leads for future investigations. Pathways “Interaction between L1-type proteins and ankyrins” and “L1CAM interactions” are dysregulated by ethanol in PD7 astrocytes (**Suppl. Figure 2**; **Figure 6A,B**, magenta arrows). L1-ankyrin signaling in FASD has been investigated in neurons at an earlier stage of development before astrocytes differentiate (46-50). L1 is a cell adhesion molecule (CAM) that exerts its adhesive properties through homophilic binding to another L1 molecule located on another cell. It is possible that astrocyte-neuron interactions are disrupted by developmental ethanol exposure through a mechanism that involves both, astrocyte and neuron L1 signaling leading to altered neuronal adhesion and development.

Several overrepresented categories of genes enriched in astrocytes and regulated (**Figure 6A**) or upregulated (**Figure 6B**) by ethanol relate to cholesterol biosynthesis (blue arrows). While in the mature brain oligodendrocytes are the major producers of cholesterol, at PD7 oligodendrocyte differentiation has not started yet and astrocytes are the major producers of cholesterol and lipoproteins that contain cholesterol (51, 52), in agreement with the observation that cholesterol biosynthetic enzymes are enriched in PD7 astrocytes. Neurons produce enough cholesterol for their survival, but cholesterol required for axonal growth and synaptogenesis is provided by surrounding astrocytes (53-55). We previously reported increased cholesterol trafficking in astrocytes *in vitro* and in the cortex *in vivo* by ethanol (56, 57). This study further supports the dysregulation of cholesterol homeostasis in FASD (58); the increased efflux of cholesterol from astrocytes and its uptake by developing neurons could be another mechanism by which ethanol increases dendritic arborization.

Pathway analysis of TRAP-seq results in non-astrocyte enriched genes revealed a strong overrepresentations of categories related to translation and ribosome biogenesis; the vast majority of these genes were downregulated (**Figure 5A,B**). Our results agree with a previous report that ribosome biogenesis and protein translation categories were overrepresented in the bulk hippocampus of mice neonatally exposed to ethanol (59) and identify protein translation as being altered by developmental ethanol exposure in astrocytes.

In conclusion, the analysis of the astrocyte translatome in a FASD mouse model of third-trimester equivalent ethanol exposure identified several novel mechanistic leads by which ethanol affects hippocampal development that are likely to permanently alter brain connectivity and be involved in some of the cognitive and behavioral effects of FASD. Categories overrepresented in pathway analyses suggest that ethanol primarily alters mechanisms of astrocyte-neuron interactions involved in neuronal development. Following up on one of these analyses, we demonstrated that ethanol inhibits the astrocyte translation of *Chpf2* and *Chsy1*, enzymes involved in the biosynthesis of inhibitory CS-GAGs, reduces the hippocampal levels of disaccharides forming CS-GAGs, and increases dendritic arborization in hippocampal pyramidal neurons. Interestingly, the silencing of *Chpf2* in astrocyte cultures results in increased hippocampal neuron neurite outgrowth. These studies provide mechanistic evidence of changes in astrocytes induced by ethanol that alter neuronal developmental trajectories and identify astrocytes and CS-GAGs as targets for FASD therapeutics. In addition to the specific mechanisms explored in this study, astrocyte translatome analysis following developmental ethanol exposure generated numerous new leads that revealed the extent of the effects of ethanol on astrocytes. The consequences of altered astrocyte functions on brain development need to be considered when studying the pathophysiology of FASD and other neurodevelopmental disorders.

## Methods and Materials

### Animals and Neonatal Alcohol Exposure

Hemizygous B6;FVB-Tg(Aldh1l1-EGFP/Rpl10a)JD130Htz/J (Aldh1l1-EGFP/Rpl10a) mice were originally created by Nathaniel Heintz (24, 25). Aldh1l1-EGFP/Rpl10a (Strain # 030247) and C57BL/6J mice were obtained from The Jackson Laboratory (JAX, Bar Harbor, ME;) and time-mated at the Portland VA. Male Aldh1l1-EGFP/Rpl10a mice were crossed with female C57BL/6J mice to generate time-pregnant litters for TRAP-seq and TRAP-qPCR experiments. Litters of Aldh1l1-EGFP/Rpl10a (TRAP experiments) or C57BL/6J (Golgi-Cox and CS-GAG experiments) mice were randomly assigned to either control (air) or ethanol vapor chambers and were exposed from PD2-PD7 between 09:00 and13:00 (4 h) each day as previously described (60). An interesting feature of this alcohol exposure paradigm is that pup blood ethanol concentrations (BECs) reach consistently high levels throughout each day of exposure (with little or no metabolic adaptation to ethanol), while dams undergo rapid metabolic adaptation and, for this reason, their BECs rapidly decrease to non-intoxicating levels and maternal care of ethanol-exposed litters is not affected (60). Time-mated Sprague-Dawley dams were purchased from Charles River. Between PD4 and PD9 Sprague-Dawley rat pups were administered ethanol via intragastric intubations (34). Rat pups that were given EtOH were also given two intubations of milk formula without EtOH at two–hour intervals starting two hours after the last EtOH intubation, to compensate for lack of suckling caused by inebriation; pups not receiving EtOH were sham-intubated at the same intervals. Intragastric intubation was done by inserting flexible tubing that was dipped into corn oil for lubrication into the esophagus of the neonatal rat as previously reported (34). Immediately following ethanol exposure, pups were euthanized, the hippocampi or the whole brains were dissected from each pup, and flash frozen in liquid nitrogen or incubated in the Golgi-Cox solutions. All animals were housed at the VA Portland Health Care System Veterinary Medical Unit, on a 12 h light/dark cycle at 22 ±1 °C. All the animal procedures were performed in accordance with the National Institute of Health Guidelines for the Care and Use of Laboratory Animals and were approved by the VA Portland Health Care System’s Institutional Animal Care and Use Committee.

### Aldh1l1-EFGP-Rpl10a mice and Translating Ribosomal Affinity Purification

Following ethanol treatments, pups were genotyped by tail biopsy for sex and presence of the transgene. Translating Ribosome Affinity Purification (TRAP) was carried out as previously described (61) on PD7 hippocampal tissue using anti-GFP antibodies (Memorial-Sloan Kettering Antibody and Bioresource Core Facility) and protein A/G magnetic beads (ThermoFisher, 88803). Following the final wash of the RNA-antibody-bead complex, RNA was extracted from TRAP and input samples with TriZol and the Direct-Zol RNA MicroPrep Kit (Zymo Research, Orange, CA).

### RNA-Seq and Analysis

TRAP and Input hippocampus samples from 12 ethanol vapor exposed (6 female, 6 male) and 12 control air exposed (6 female, 6 male) mice for a total of 48 samples were sequenced at the Integrated Genomics Lab core facility at OHSU. RNA quality was assessed by Agilent Bioanalyzer; all sequenced samples had a RIN ≥ 8.8 (**Suppl. Table 1**). RNA-Seq libraries were profiled on a 4200 Tapestation (Agilent) and sequenced on a NovaSeq 6000 (Illumina). Before sequencing, TRAP samples were tested for enrichment of astrocyte markers and depletion of neuronal, oligodendrocyte, and microglia markers in comparison to the respective input samples by qPCR. Primers for mouse *Aldh1l1* (Forward: 5’-GGTTTACTGCCAGCTAAGGAAG-3’; Reverse: 5’-CACGTTGAGTTCTGCACCCA-3’), *S100b* (Qiagen QT00151536), *Tubb3* (Forward: 5’-CCCAGCGGCAACTATGTAGG-3’; Reverse: 5’-CCAGACCGAACACTGTCCA-3’), *Rbfox3* (NeuN) (Forward: 5’-GGCAAATGTTCGGGCAATTC-3’; Reverse: 5’-TCAATTTTCCGTCCCTCTACGAT-3’), *Cd11b* (Itgam) (Forward: 5’-GGCTCCGGTAGCATCAACAA-3’; Reverse: 5’-ATCTTGGGCTAGGGTTTCTCT-3’), *Mbp* (Forward: 5’-GGCAAGGTACCCTGGCTAAA-3’; Reverse: 5’-GTTTTCATCTTGGGTCCGGC-3’), and *Opalin* (Forward: 5’-CAAGGGGGCCTGTCAGAAAT-3’; Reverse: 5’-GCTGCCAGTCCAATAGAGGG-3’) were used with the Luna Universal One-Step RT-qPCR Kit (NEB) with SYBR Green detection on the CFX96 (Bio-Rad) thermocycler. Cycle threshold values were normalized to total RNA using RiboGreen (ThermoFisher) and expressed as log2 transformed data relative to the input samples. Fastq files were assembled from the base call files using bcl2fastq (Illumina). The fastq files were trimmed with Trimmomatic (62) using the built-in filters for Illumina adapters. After trimming, sequence files were aligned with STAR aligner (63). The reference genome was Mus musculus GRCm38, downloaded with annotations from Ensembl. Following the alignment, the SAM files were converted to BAM format using SAMtools (64). Raw read count data were analyzed using the R package DESeq2 (65). Genes with less than a total of 10 reads across the 48 samples were excluded from the analysis. Clustering and principal component analysis was conducted on all 48 samples (inputs and IP) as well as analysis of differential expression between the IP and input fractions. Differential expression/translation analysis by ethanol treatment was carried out in each fraction separately. Significance of RNA-Seq data was determined by DESeq2 false discovery rate (FDR) adjusted Wald test p-values less than 0.05. Raw and processed RNA-Seq data are available at the NCBI Gene Expression Omnibus (https://www.ncbi.nlm.nih.gov/geo/) accession number GSE272272.

### Bioinformatic analysis

Gene Ontology (GO) and pathway enrichment analysis was carried out using the enrichR package in R to query the enrichr database (66, 67). We limited the analysis to the following databases to focus our results on GO categories and known gene pathways: “GO_Molecular_Function_2021”, “GO_Cellular_Component_2021”, “GO_Biological_Process_2021”, “BioPlanet_2019”, “Elsevier_Pathway_Collection”, “KEGG_2021_Human”, “MSigDB_Hallmark_2020”, “WikiPathway_2021_Human”.

### Golgi-Cox staining and morphometric analysis

Twenty-eight C57BL/6J mice (16 female) from 9-10 litters that were exposed to ethanol vapor and 9 litters that were exposed to air between PD2 and PD7 were used in these studies. Brains were stained using the FD Rapid GolgiStain Kit (FD Neuro-Technologies Inc., Columbia, MD) according to the manufacturer’s instructions and as previously described (34, 68) with minor modifications made to optimize the method for use in neonatal mouse brains. Briefly, brains were immersed in the Golgi-Cox impregnation solution and stored at room temperature for 3 weeks in the dark and then transferred to Solution C and stored at 4° C for 2 weeks. The brains were then rapidly frozen and embedded in Tissue Freezing Medium before sectioning on the cryostat. Coronal 100 μm sections were mounted on gelatin-coated microscope slides, dried at room temperature overnight, and stained the following day. Image stacks were collected with a Leica DFC365 FX camera (Exposure = 6.0ms, Binning =1x1). Pyramidal neurons were imaged and the complete dendritic tree was traced with the software Neurolucida 360 (MBF Bioscience, Inc.). Eight cells/brain (4 cells/hemisphere) were imaged using a 40x objective. Only fully impregnated CA1 pyramidal neurons fully present in a single section and clearly distinguishable from neighboring neurons were measured. For apical dendrites, we analyzed the following measures: *complexity*, defined as [Sum of Terminal Orders + Number of Ends] * [Total Dendritic Length/Number of Dendrites]; *sum of the terminal orders*, defined as the number of “sister” branches encountered as you proceed from the terminal to the cell body and calculated per each terminal; *number of ends*; *sum of branch orders*, defined as the sum of the terminal order plus the number of ends; *total dendrite length* (in μm); *maximal terminal distance*, defined as linear distance of the furthest dendrite from the cell body. For basal dendrites, in addition to the measures analyzed for apical dendrites, we also analyzed *number of basal dendrites per cell*; the length of basal dendrites was analyzed in two ways: as *average dendrite length* per cell (because pyramidal cells usually have more than one basal dendrite) and as *total dendrite length*. Because the basal and the apical dendritic trees of pyramidal neurons are morphologically different, we analyzed apical and basal dendrites separately.

### Spine Quantification

Spine density analyses were performed on neurons previously utilized for dendritic arborization measures. Seven new (not previously analyzed) neurons were analyzed in order to ensure a minimum of 500 µm of dendrite per group, and at least 200 spines per group (69, 70). Only secondary apical dendrites, identified by shaft ordering, were chosen for analysis. Basilar processes were not included due to limited segments meeting selection length criteria. Spines were imaged using a Leica DM500b microscope with 100x oil immersion objective. Image stacks (z-section thickness = 0.1 µm) between 5 and 20µm thick were collected with a Leica DFC365 FX camera (Exposure = 6.0ms, Binning =1x1). Image stacks were reconstructed to 3D volumes using Neurolucida 360 (MBF Bioscience, Inc.). Dendritic branches were traced with the Neurolucida contour function through the z-plane to obtain accurate length measures for selection criteria. Images of dendritic sections were collected, excluding the last (distal) 10µm of the branch. Due to the fragility of the neonatal tissue, branches with minor (<3 µm) breaks were included in analysis. Neurons were excluded if they did not have a 20 µm or longer secondary apical dendrite, had >3 µm breaks in the branch, or if the analyzed dendrite was grossly overlapped by neighboring branches. After tracing the dendritic region of interest with the directional kernel based semi-manual or smart manual tracing function (Fig.1D), spines were semi-manually identified using the Neurolucida spine identification function (outer range = 2.5 µm, minimum height 0.3 µm, detector sensitivity = 100%, minimum count = 10 voxels) by an experimenter blind to both treatment condition and sex (71). A single average spine density value was calculated for each animal from up to 8 dendritic segments (10 to 40 µm length) from 2-4 pyramidal neurons. In total, 1318 µm and 1108 µm of dendritic branch was analyzed for control and treatment animals, respectively. Spine density was calculated as the number of spines per length of dendrite in µm as well as by the number of spines per length of dendrite in µm per diameter of dendrite in µm to account for changes in surface area (width corrected).

### RNA-Fluorescence In Situ Hybridization (FISH)

Naïve PD7 mouse brains were dissected and flash frozen in isopentane chilled in a dry-ice isopropanol bath then stored at -80° C until sectioning. Fresh frozen brains were sectioned on a Leica CM3050S Cryostat at 100 µm and collected on Superfrost plus slides. HCR^TM^ RNA-FISH for *Aldh1l1*, *Tubb3*, and *Chpf2* or *Chsy1* was carried out according to the manufacturer’s recommendations with the omission of the optional Proteinase K step and nuclear staining with DAPI (Molecular Instruments; Los Angeles, CA, USA). RNA-FISH was imaged using a Leica DM500b microscope with a DFC365 FX camera.

### GAG Disaccharide Analysis by Liquid Chromatography/Mass Spectrometry (LC/MS) and Quantification of CS-GAG Disaccharide Concentrations

CS-GAG disaccharide levels were analyzed in the hippocampus of control and ethanol-exposed rat pups as previously described (42, 68, 72). Samples were first defatted by the treatment with 0.2 mL acetone for 30 min, vortexed, and the acetone supernatants were discarded. The remaining tissue samples were dried in the hood, lyophilized, and subjected to proteolysis at 55 °C with 10% actinase E (10 mg/mL) until all tissue was dissolved. GAGs were purified by Mini Q spin columns. Samples eluted from Mini Q spin column were desalted by passing through a 3 kDa molecular weight cut off spin column and washed three times with distilled water. The casing tubes were replaced before 150 μL of digestion buffer (50 mM ammonium acetate containing 2 mM calcium chloride adjusted to pH 7.0) was added to the filter unit. Recombinant heparin lyase I, II, III (pH optima 7.0–7.5) and recombinant chondroitin lyase ABC (10 mU each, pH optimum 7.4) were added to each sample and mixed well. The samples were all placed in a water bath at 37 °C for 12 h, after which enzymatic digestion was terminated by removing the enzymes by centrifugation. The filter unit was washed twice with 300 μL of distilled water and the filtrates containing the disaccharide products were dried via vacuum centrifuge. Half of the dried samples were 2-aminoacridone (AMAC)-labeled by adding 10 μL of 0.1 M AMAC in dimethylsulfoxide/acetic acid (17/3, V/V) and incubated at room temperature for 10 min, followed by addition of 10 μL of 1 M aqueous sodium cyanoborohydride and incubation for 1 h at 45 °C. A mixture containing all 17-disaccharide standards prepared at 0.5 ng/μL was similarly AMAC-labeled and used for each run as an external standard. After the AMAC-labeling reaction, the samples were centrifuged, and each supernatant was recovered. LC was performed on an Agilent 1200 LC system at 45 °C using an Agilent Poroshell 120 ECC18 (2.7 μm, 3.0 × 50 mm) column. The mobile phase A (MPA) was a 50 mM ammonium acetate aqueous solution, and the mobile phase B (MPB) was methanol. The mobile phase passed through the column at a flow rate of 300 μL/min. The gradient was 0–10 min, 5–45% B; 10–10.2 min, 45–100%B; 10.2–14 min, 100%B; 14–22 min, 100–5%B. Injection volume was 5 μL. A triple quadrupole mass spectrometry system equipped with an ESI source (Thermo Fisher Scientific, San Jose, CA) was used as detector. Disaccharide quantification was performed by comparing the integrated disaccharides peak area with the external disaccharide standard peak area. The data analysis was performed in Thermo Xcalibur software. The final GAG disaccharide concentrations were calculated by dividing the GAG disaccharides mass per sample (ng) by the wet weight of each hippocampal sample and expressed as ng/mg tissue.

### Astrocyte and neuron primary culture, astrocyte Chpf2 silencing, and neurite outgrowth measurements

Astrocyte cultures were prepared from gestational day (GD) 21 fetuses, as previously described (61, 73, 74). Astrocytes were maintained in Dulbecco’s Modified Eagle’s Medium (DMEM; Thermo-Fisher Scientific), supplemented with 10% Fetal Bovine Serum (Atlanta Biologicals, Flowery Branch, GA), 1000 units/mL penicillin–streptomycin (pen-strep, Thermo-Fisher Scientific) in a humidified incubator at 37 °C under a 5% CO_2_/95% air atmosphere until confluent (7-10 days *in vitro*). Astrocytes were then sub-cultured on 6-well plates at the density of 0.4x10^6^ cells/well. Two days after sub-culture, astrocytes were transfected in DMEM/0.1% BSA without antibiotics supplemented with 0.5 ml/well Lipofectamine RNAiMAX (Thermo-Fisher Scientific) diluted with Opti-MEM I Reduced Serum Media (Opti-MEM; 1:50 dilutions; Thermo-Fisher Scientific) containing 1 nM Stealth siRNA targeting the rat Chpf2 (Invitrogen/Thermo-Fisher Scientific, Carlsbad, MA) or 1 nM Stealth siRNA non-target control, medium guanosine and cytosine (GC) content (non-target siRNA; Invitrogen/Thermo-Fisher Scientific) for 24 h. Transfection medium was then replaced with DMEM/0.1% BSA/pen-strep for additional 48 h. This procedure is similar to the one previously published by us to silence other proteins (61, 74). The astrocyte conditioned medium (ACM) from control and *Chpf2*-silenced astrocytes was collected and centrifuged at 200 x *g* for 10min to eliminate cell debris. Hippocampal neuron cultures were prepared from fetuses at GD17-18 as previously described (73, 74) and plated on poly-D-lysine-coated glass coverslips in complete Neurobasal medium containing B27 supplements at the density of 1 x 10^4^ neurons/coverslip. After 4h the medium was replaced with ACM medium prepared from control and Chpf2-SiRNA-treated astrocytes for an additional 48 h. At the end of the incubation, neurons were fixed in 4% paraformaldehyde and immunolabeled with a neuronal-specific anti-β-III tubulin antibody (1:150 dilution; Millipore, Burlington, MA, #MAB1637), followed by a goat anti-mouse Alexa Fluor 488 secondary antibody (1:300 dilution; Thermo-Fisher Scientific, #A11001), as previously described (61, 73, 74). Glass coverslips were then mounted on microscope slides. β-III tubulin-labeled neurons were imaged on a Leica DM5000b microscope attached to a DFC365 FX camera. Pictures of neurons obtained with the 40x objective were traced and analyzed with Neurolucida 360 by a researcher blind to the experimental treatments. 29-30 neurons/coverslip from 3-4 coverslips/treatment were selected based on the following parameters: 1) had three or more neurites that were all longer than the cell body (pyramidal neurons), 2) did not overlap with other neurons, 3) had no breakage in the cell bodies or neurites, and 4) were surrounded by a similar number of neurons (fields that at 10x magnification show only sparse neurons, 1 or 2 per field, were not considered for analysis).

### Statical Analyses

Neuronal morphology, GAGs levels, and neurite measurements were analyzed using a linear mixed effects analysis in R (75) using the package lme4 (76) as previously described (34). For neuronal morphology analyses, fixed effects were animal and litter to account for multiple neurons measured per animal and multiple animals per litter. For GAGs analysis, litter was entered as a fixed effect to account for multiple animals analyzed from each litter. For neurite analysis, coverslip was a fixed effect to account for multiple neurons measured for each coverslip. qRT-PCR results were analyzed by a two-way ANOVA.

## Acknowledgements

This study was supported by NIH P60AA010760, R01AA029486, and U01AA029965, and VA Merit Review Award I01BX001819, and by facilities and resources at the Portland VA Health Care System to MG. The contents of this article do not represent the views of the United States Department of Veterans Affairs or the United States government. We thank Ms. Venice Loar for her help in the neurite outgrowth measurements, Ms. Natalie Gorham for her help in generating the RNA-FISH images, and Dr. Deborah Finn and Ms. Melinda Helms for their help with blood ethanol concentration determinations.

---

**Supplemental Figure 1.**
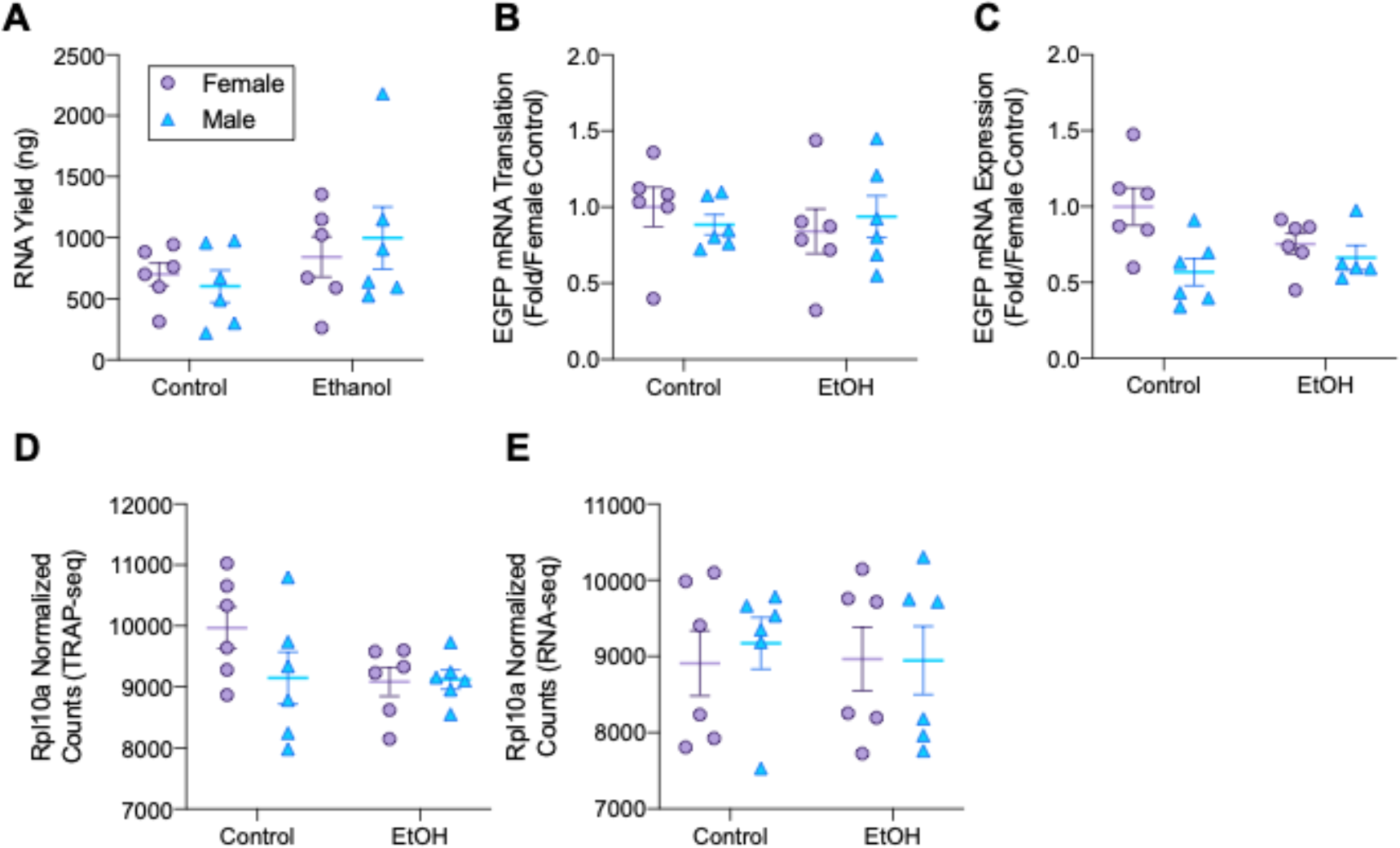
**A**: RNA yield from TRAP samples was not different across sex or treatment. **B**: EGFP mRNA translation in TRAP samples showed no sex or treatment effects by qRT-PCR. **C**: EGFP mRNA in input samples showed no sex or treatment effects by qRT-PCR. **D**: Rpl10a normalized counts in TRAP-Seq samples showed no significant differences by treatment or sex. **E**: Rpl10a normalized counts in RNA-Seq (input) samples showed no significant differences by treatment or sex.

**Supplemental Figure 2.**
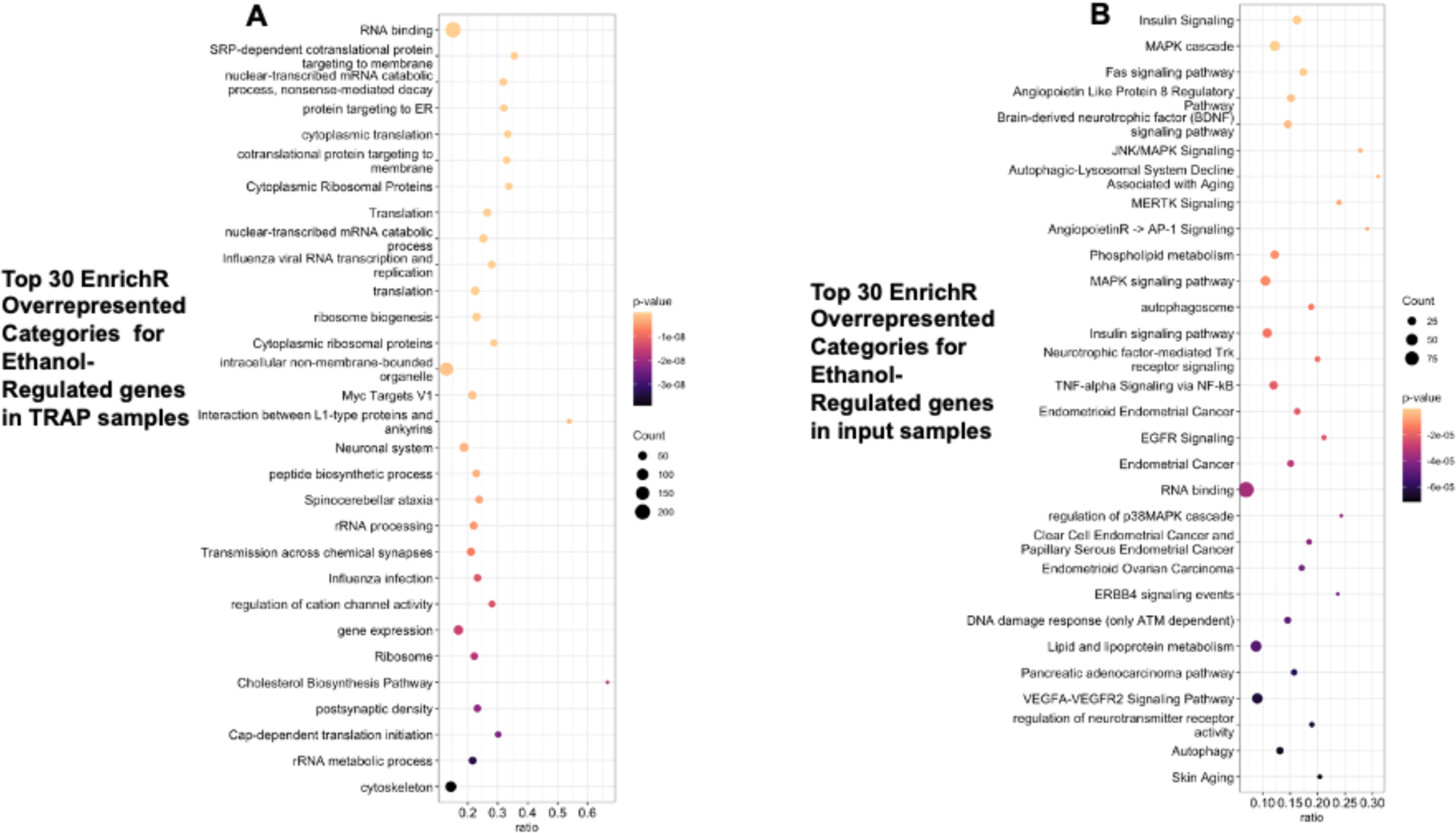
Gene category overrepresentation summary plots for ethanol regulated genes. **A:** Top 30 EnrichR categories for genes that were regulated in TRAP samples. **B**: Top 30 statistically significant Enrichr overrepresented categories for genes that were regulated in input samples.

**Supplemental Figure 3.**
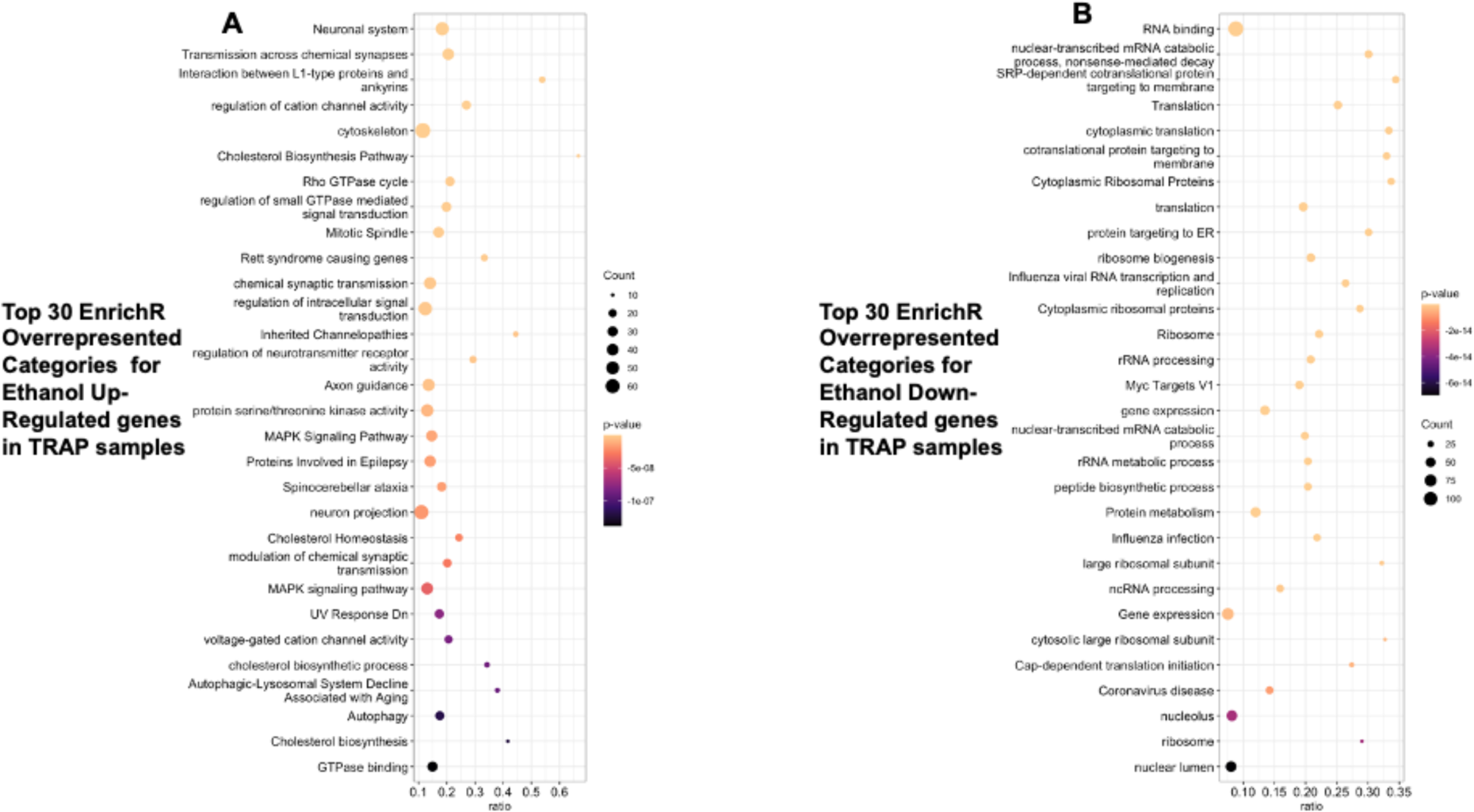
Gene category overrepresentation summary plots for ethanol regulated genes in the TRAP fraction. **A:** Top 30 Enrichr overrepresented categories for genes that were upregulated by ethanol. **B**: Top 30 Enrichr overrepresented categories for genes that were downregulated by ethanol.

**Supplemental Figure 4.**
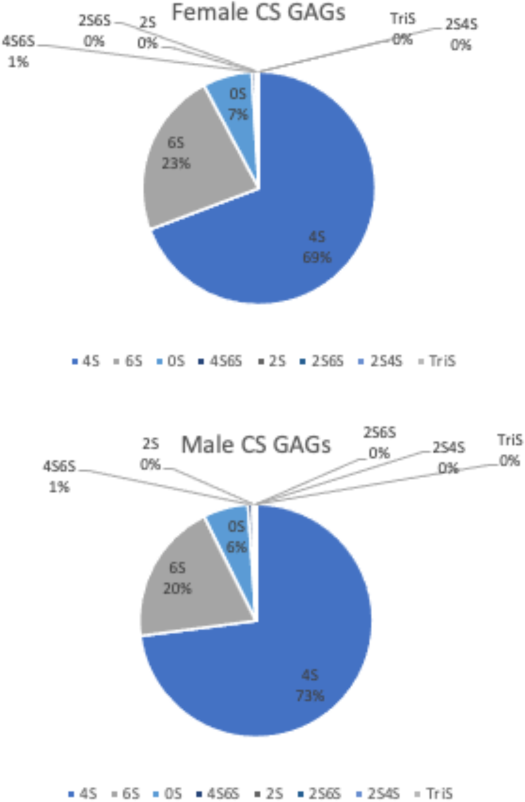
Relative distribution (expressed as percentage of total CS-GAGs) of CS-2S4S6S (TriS); CS-2S4S; CS-2S6S; CS-4S6S; CS-2S; CS-4S; CS-6S; CS-0S disaccharides in the neonatal hippocampus of female (top) and male (bottom) rat pups.

